# Evolution of tandem repeats in putative CSP to enhance its function: A recent and exclusive event in Plasmodium vivax in India

**DOI:** 10.1101/2023.11.28.568961

**Authors:** Manoswini Dash, Veena Pande, Aparup Das, Abhinav Sinha

## Abstract

The molecular hitchhiking model proposes that linked non-coding regions also undergo fixation, while fixing a beneficial allele in a population. This concept can be applied to identify loci with functional and evolutionary significance. Putative circumsporozoite protein (CSP) in Plasmodium vivax (PvpuCSP) identified following the molecular hitchhiking model, holds evolutionary significance. We investigated the extent of genetic polymorphism in PvpuCSP and the role of natural selection which shapes the genetic composition and maintains the diversity in *P. vivax* isolates from India. Sequencing the putative CSP of *P. vivax* (PvpuCSP) in 71 isolates revealed a well-conserved N- and C-terminal, constituting around 80% of the gene. PCR amplification and sequencing validated extensive diversity in the repeat region, ranging from 1.8 to 2.2 kb towards the C-terminal, identifying 37 different alleles from 71 samples. The recent and exclusive accumulation of repeats in puCSP within *P. vivax* highlights its highly variable length polymorphism, making it a potential marker for estimating diversity and infection complexity. Episodic diversifying selection in the PvpuCSP repeat region, evidenced by statistically significant p-values and likelihood ratios, enhances amino acid diversity at various phylogenetic levels, facilitating adaptation for accommodating different substrates for degradation.

## Background

India is endemic to malaria mainly with two Plasmodium species, *vivax* and *falciparum*, equally prevalent with sporadic geographic distribution. Of them, *P. vivax* remained neglected for ages owing to its non-fatal attribute and at present a major hindrance in the malaria elimination program (Habtamu, Petros, and Yan 2022). However, reports of severity caused by *P. vivax* from many parts of the world have changed the assumption of benign nature of the *P. vivax* infection. When malaria elimination is the goal to achieve by many developing countries in the near future, it is of critical importance to equally target *P. vivax* to unfold the genetic features, gain a clear insight into its diversity and distribution in the population, for the effective design and implementation of control strategies.

Syntenic chromosomal segments are regions of chromosomes where the gene, gene order, and orientation are similar between two species (Tang et al. 2008). Conserved syntenic regions represent common ancestry and a measure of divergence among species. Genomic research has largely exploited the “syntenic feature” of the genome to improve the mapping accuracy of a newly sequenced genome and identify conserved non-coding sequences. Comparative genomic studies have revealed many syntenic chromosomal segments between *P. falciparum* and *P. vivax* genomes (Carlton, Adams, et al. 2008; Carlton, Escalante, et al. 2008; Gupta et al. 2010a). Population genetic analysis of one such 200 kb long syntenic region revealed a substantial decline in genetic diversity in the non-coding (intronic and intergenic) regions, suggesting a recent event of molecular hitchhiking (Gupta et al. 2010b; Gupta, Srivastava, and Das 2012). Genetic hitchhiking, also known as the selective sweep, is a phenomenon in which the genetic diversity of a linked allele is lost in the process of establishing a beneficial trait in the population (Barton 2000; Maynard Smith and Haigh 2008). In other words, investigating non-coding loci with high levels of genetic homogeneity can help identify the nearest beneficial trait under selection.

The locus named circumsporozoite protein (*PV*X_086150; further referred as *Pv*puCSP) is adjacent to a non-coding stretch with reduced diversity within the syntenic region, suggesting that it may be under selection pressure (Gupta et al. 2012). This locus is named after the circumsporozoite protein, which is a vaccine target for *P. falciparum*. Our interest in *Pv*puCSP is due to its evolutionary significance and its potential role in the *P. vivax* life cycle. However, there is currently no experimental evidence to support its cellular localization, stage-specificity, or role in the *P. vivax* life cycle. A systematic and thorough computational study conducted by our team revealed some exciting and distinguishing features about the locus (Dash, Pande, and Sinha 2019). The locus has a domain organization for E3 ubiquitin ligase, and the protein product participates in the ubiquitination pathway. The study provided strong computational evidence to suggest that the locus (*PV*X_086150) is different from the known vaccine candidate (*PV*X_119355). To our interest, the intra-genic pattern of *Pv*puCSP has a strong resemblance to the well-known CSP of *P. falciparum* in terms of a highly conserved N and C-terminal domain and a hyper-variable central repeat region (Dash et al. 2019). Hence, we aimed to analyse the genetic structure, estimate diversity, and assess distinctiveness of the locus in *P. falciparum* and *P. vivax*. Additionally, we sought to infer the impact of natural selection on the locus for a deeper understanding of its evolutionary significance.

## Results

The nested PCR assay revealed 90 mono-*P. vivax* infection out of 158 samples tested (Supplementary Table 3). The *Plasmodium* infection was confirmed by a 1100 bp band in the first step of nested PCR (Fig 1-a), and the species was detected by two bands at 205 bp and 120 bp in the second step for *P. falciparum* and *P. vivax*, respectively (Fig 1-b). Samples co-infected with *P. falciparum* and *P. vivax* (further used as mixed infection) has shown two corresponding bands. The *Pv*puCSP gene was sequenced only in samples infected with *P. vivax* (n=90) (Supplementary Table 3). The *Pv*puCSP exhibits a highly variable repeat region at its C-terminus, as observed in gel electrophoresis with bands of varying lengths (Fig 1-c). The study used the entire *Pv*puCSP gene from 64 isolates. Partially sequenced genes were excluded, except for the isolates with sequenced repeat region, used for VNTR analysis.

**Figure 1:**
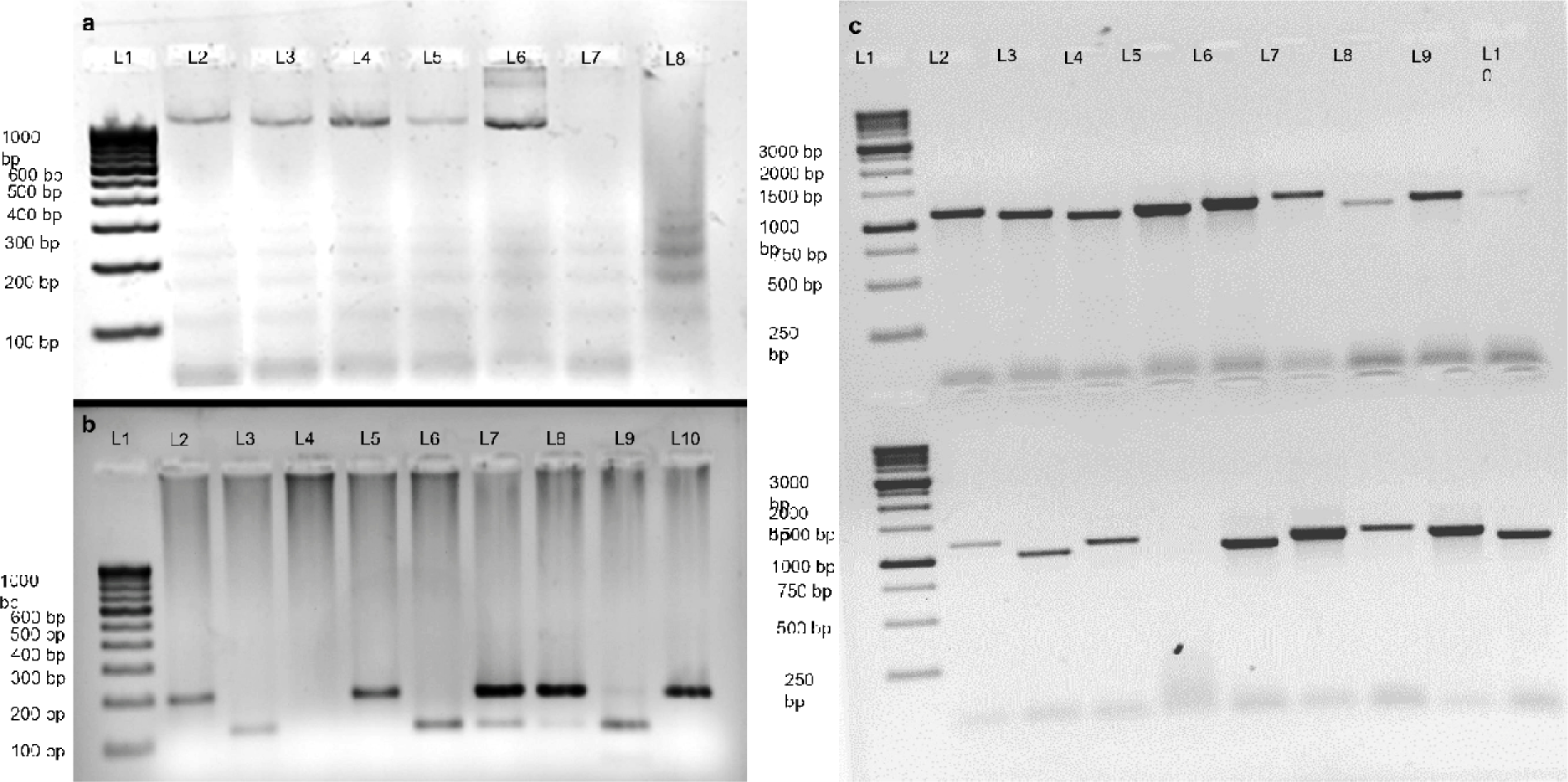
Identification of *Plasmodium* species and presence of variable repeat region in *Pv*puCSP using nested PCR assay. (a) The first step of nested PCR confirms the presence of *Plasmodium* with the band around 1100 bp. L1: 100 bp DNA marker, L2-L8: representative samples. L2 to L6: positive for *Plasmodium* infection, L7 and L8: *Plasmodium* negative (b) Second step of nested PCR confirms the species *P. falciparum* and *P. vivax*. L2, L5, and L10: mono-Pf positive, L3, L6 and L9: mono-*Pv* positive, L7 and L8: mixed infection, L4: Negative. (c) Presence of repeat region in *Pv*puCSP was confirmed by amplifying the repeat region (1168 bp) showing band of varying length. Representative *P. vivax* samples were amplified using a mentioned set of primers. Note: The amplicon lengths are determined based on the puCSP of the reference strain *P. vivax* Sal-1

### *Pv*puCSP has a distinctive hyper-variable repeat region

The *Pv*puCSP analysis by PCR assay and tandem repeat analysis of confirmed the presence of hyper-variable repeat regions (1.8-2.2 kb) towards the C-terminal (Fig 2-a; Supplementary Table 4). The consensus of the repeat is AGG[GA/TG]TAA[C/T]GC[C/T], where 4^th^, 5^th^, 9^th^, and 12^th^ positions are varying thereby generated four RATs that code for two PRMs i.e. Arg-Asp-Asn-ala (R-D-N-A) and Arg-Cys-Asn-ala (R-C-N-A) (Fig 2-b). The R-D-N-A is coded by 3 RATs (AGGGATAACGCC [R1], GGGATAACGCT [R2], AGGGATAATGCC [R3]) and R-C-N-A has single RAT (AGGTGTAATGCC [R4]) (Fig 2-d). The variation in the PRM is due to change in the second codon of the repeating unit from Asp ➔ Cys. Alleles were classified based on the arrangement of RATs in repeat regions (Supplementary Table 2; Fig 2-c). In most of the isolates, the repeats were started with R1 and ended with R3, except in three isolates where the first repeat is R2 (G3948-GUJ and K13-ODI) and the last repeat is R1 (D2115-DEL). The range of number of repeat units varies from 2 to 30, generating a length variation ranging from 24 to 360 bp. The repeat region of *Pv*puCSP was sequenced in 71 *P. vivax* isolates and included 889 individual tetra-peptide repeats, of which 92% (817) encoded RDNA (R1: 33%, R2: 41%, R3:7%) while rest 8% (72) encoded RCNA.

**Figure 2:**
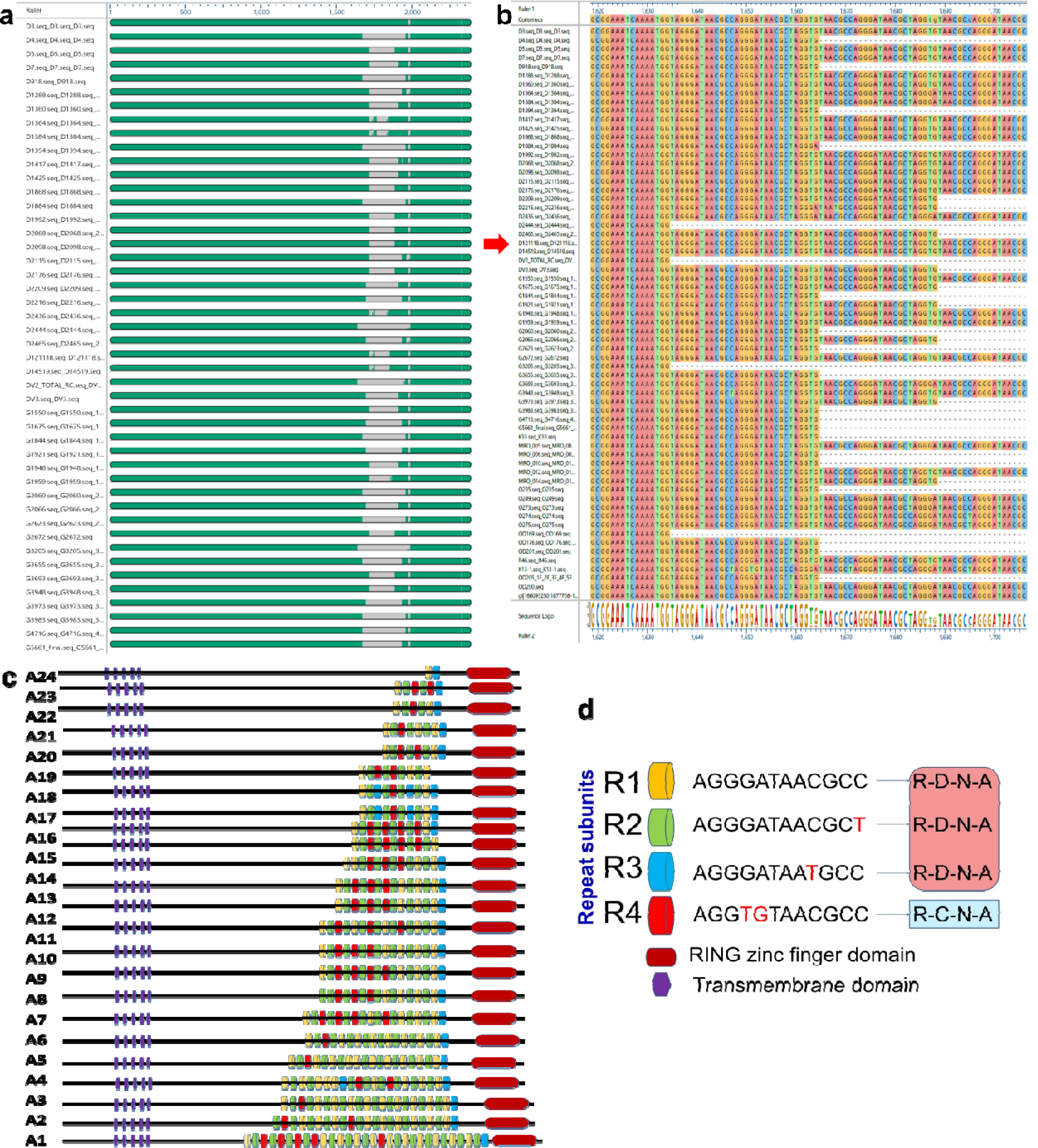
MSA of *Pv*puCSP gene sequenced in 64 isolates showing the repeat region and pattern of repeats. (a) MSA of complete *Pv*puCSP gene, grey space shows the gaps. (b) MSA showing the repeat region of *Pv*puCSP. (c) Alleles were formed based on the arrangement pattern of repeat allotypes. The transmembrane domain and RING zinc finger domain were shown as a purple rectangle and red cylindrical shape, respectively(d) Arrangement of nucleotides to form different RAT and PRM in the repeat region.

### *Pv*puCSP has highly diverse repeat regions in Odisha

Based on the repeats, a total of 37 alleles/haplotypes were established from the 71 isolates (Fig 3-b), with 18 haplotypes from Delhi and Odisha and 13 haplotypes from Gujarat (Supplementary Table 6. Haplotypes varied across populations, mostly being population-specific (Fig 3-a). Odisha had the highest exclusive haplotypes (n=13), followed by Delhi (n=12) and Gujarat (n=7 Five haplotypes (A12, A15, A24, A25, and A27) were shared among all populations, and one haplotype (A10) was shared between Delhi-Gujarat and Gujarat-Odisha (A23) populations (Fig 3-a). Despite an equal number of haplotypes in Delhi and Odisha, Odisha had a higher effective number of alleles (Ne) due to a higher frequency of alleles, while Delhi had mostly rare haplotypes (Fig 3-c). The overall haplotype diversity was higher (>0.5) in each population, with Odisha exhibiting a higher unbiased haplotype diversity (He=0.975). Shannon’s information index (I), expected heterozygosity, and allelic richness (Fig 3-c; Supplementary Table 4). PRM-based analysis supported this, showing high genetic diversity in Odisha, followed by Delhi and Gujarat, with alleles distributed almost equally in Gujarat (Fig 3-a).

**Figure 3:**
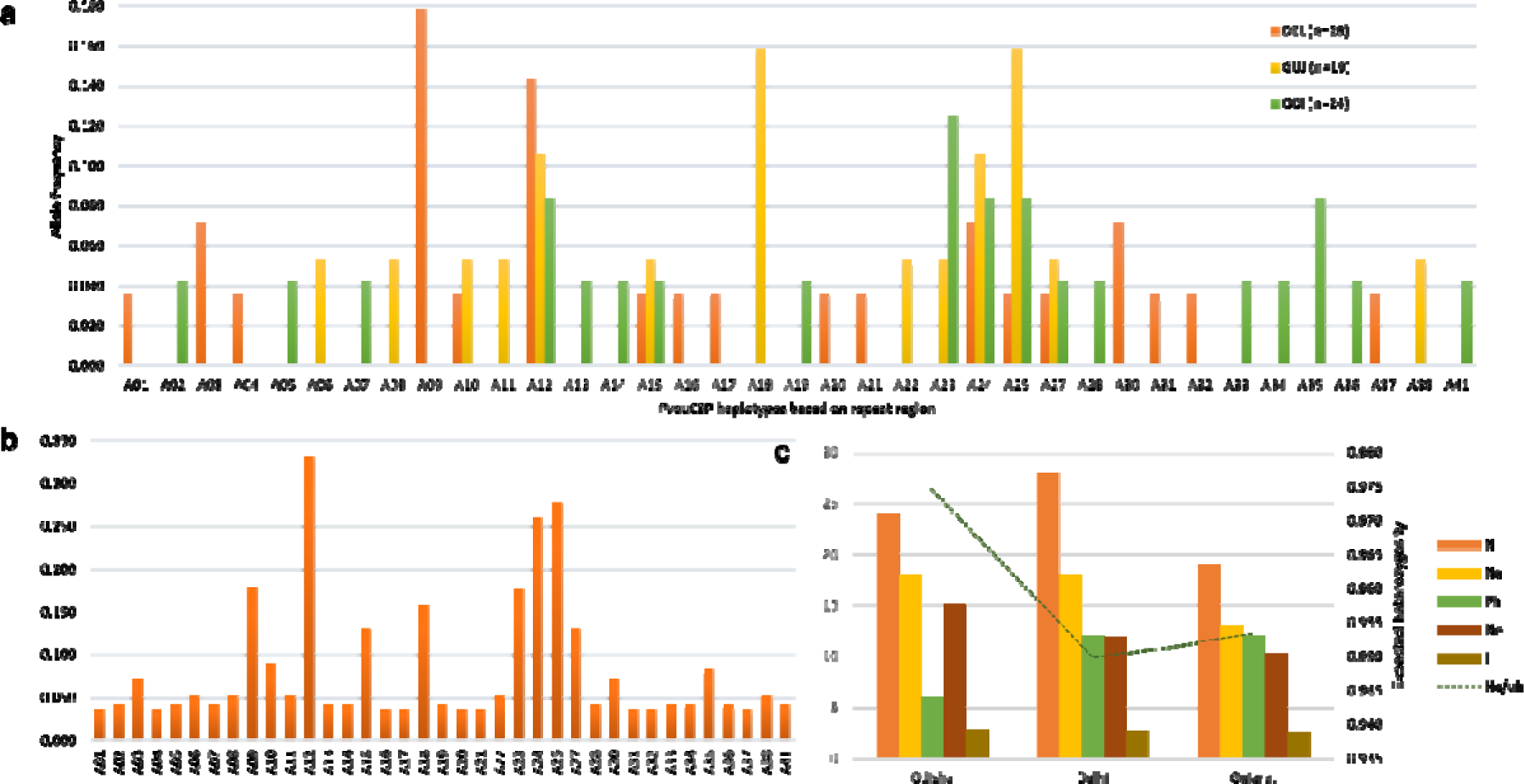
Distribution and frequency of haplotypes based on repeat region of *Pv*puCSP. (a) Population-wise distribution of the alleles showing the common and unique alleles in each population (b) The total alleles and their frequency distribution in total samples (c) Haplotype diversity and their frequency distribution of based on arrangement of repeat allotypes (RATs) in the repeat region of *Pv*puCSP in Odisha, Delhi and Gujarat populations. H_d_: Haplotype Diversity, N_a_: Number of haplotypes, I: Shannon’s Information Index, N_e_: Effective number of alleles.

### *Pv*puCSP, except the repeat region, is less polymorphic

The SNP analysis was performed excluding the repeat region. There were 8 polymorphic sites reported from the non-repeat region, of which 4 are parsimony informative sites (98, 387, 1643, and 1749) and four singletons (1235, 1493,1978, and 1989). Out of total SNPs, three were synonymous (aa position: 129, 583 and 663) and five were non-synonymous (Pro33Gln, Arg412Leu, Arg498Leu, His548Pro, Asp660His) sites (Supplementary Table 7). Two synonymous SNPs at amino acid positions 129 and 583 were found in all the three populations while all SNPs reported were present in Odisha population.

Based on the SNPs in *Pv*puCSP, there were 11 unique haplotypes (H) in 64 isolates, with haplotype diversity (H_d_) 0.683. Odisha harbors highest haplotypes (H: 9 and H_d_:0.758) followed by Delhi (H: 8 and H_d_: 0.706) and Gujarat with minimum haplotypes (H: 3 and H_d_: 0.542). The two other genetic diversity estimators (π and θ) were also implemented for each population as well as for total samples. The overall nucleotide diversity of *Pv*puCSP (π= 0.00047, θ_W_=0.00084) was less in comparison to its repeat region (π= 0.00513, θ_W_=0.01762). The diversity of repeat region was estimated by estimating nucleotide diversity of gaps/missing data, since the gaps generated here were not due to any sequencing artifact but tandem repeats. The C-terminal region and repeat region have higher genetic diversity in Odisha followed by Delhi and Gujarat. However, the N-terminal showed higher diversity in Delhi followed by Gujarat and Odisha. The analysis revealed that the overall diversity that exists in *Pv*puCSP locus was due to the extreme variation in repeat regions while other parts of the gene are relatively conserved (Table 2).

**Table 2:** Summary statistics of nucleotide variation of *Pv*puCSP in three population samples of *P. vivax*.

| Population | n | sites | S | H | H<sub>d</sub> | $\pi$ | $\theta_w$ | Tajima's D | Fu & Li's D* | Fu & Li's F* | Fu's F |
| --- | --- | --- | --- | --- | --- | --- | --- | --- | --- | --- | --- |
| AllSeq | 64 | 2013 | 8 | 1<br>1 | 0.683 | 0.00047 | 0.00084 | -1.15787 | -1.70803 | -1.71147 | -6.546 |
| Delhi | 28 | 2013 | 6 | 8 | 0.706 | 0.00048 | 0.00064 | -0.67305 | -0.57646 | -0.65106 | -4.246 |
| Gujarat | 18 | 2013 | 3 | 3 | 0.542 | 0.00029 | 0.00029 | 0.00096 | -0.5522 | -0.46362 | -0.005 |
| Odisha | 18 | 2013 | 5 | 9 | 0.758 | 0.0006 | 0.00101 | -1.36637 | -1.72671 | -1.8746 | -6.094 |
| N-Terminal (Delhi) | 28 | 1635 | 3 | 4 | 0.611 | 0.00043 | 0.00047 | -0.2093 | -0.24045 | -0.26789 | -0.491 |
| N-Terminal (Gujarat) | 18 | 1635 | 1 | 2 | 0.471 | 0.00029 | 0.00018 | 1.16615 | 0.66689 | 0.83412 | 1.215 |
| N-Terminal (Odisha) | 18 | 1635 | 2 | 3 | 0.386 | 0.00025 | 0.00036 | -0.73968 | -0.5522 | -0.6314 | -0.675 |
| N-Terminal (1-1635) | 64 | 1635 | 4 | 5 | 0.527 | 0.00036 | 0.00052 | -0.65251 | -1.30168 | -1.28604 | -1.34 |
| C-Terminal (Delhi) | 28 | 378 | 2 | 3 | 0.262 | 0.00071 | 0.00136 | -0.99368 | -0.71444 | -0.91462 | -1.128 |
| C-Terminal (Gujarat) | 18 | 378 | 1 | 2 | 0.111 | 0.00029 | 0.00077 | -1.16467 | -1.49949 | -1.61172 | -0.794 |
| C-Terminal (Odisha) | 18 | 378 | 4 | 5 | 0.601 | 0.00185 | 0.00308 | -1.19565 | -1.6133 | -1.72277 | -0.377 |
| C-terminal (1996-2376) | 64 | 378 | 4 | 5 | 0.33 | 0.00093 | 0.00225 | -1.26019 | -1.30168 | -1.43604 | -2.195 |
| Repeat Region (Delhi) | 28 | 360 | 1 | 2 | 0.138 | 0.00573 | 0.01071 | -0.741 | 0.8153 | 0.3362 | -0.38 |
| Repeat Region (Gujarat) | 18 | 360 | 1 | 2 | 0.111 | 0.00463 | 0.01211 | -1.1647 | 0.8846 | 0.4898 | -0.794 |
| Repeat Region (Odisha) | 18 | 360 | 1 | 2 | 0.111 | 0.00463 | 0.01211 | -1.1647 | 0.8846 | 0.4898 | -0.794 |
| Repeat Region (1636-1995) | 64 | 360 | 2 | 3 | 0.121 | 0.00513 | 0.01762 | -1.1948 | 0.7234 | 0.1632 | -2.137 |

#### Neutrality could not be rejected based on frequency-based tests

Tajima’s D, Fu and Li’s D, F, D* and F*, Fu’s F and Fay and Wu’s H, and Fu’s F and Fay and Wu’s H tests were performed on the total samples and sub-groups bases on populations (Delhi, Gujarat, and Odisha) and domains (amino terminal, repeat region and carboxy-terminal With the exception of Gujarat and Delhi, there was a tendency towards departure from neutrality in all categorized analyses, although the deviations were statistically non-significant (Table 2). Negative values for most estimators across all populations indicate that pairwise nucleotide differences are less than segregating sites. Tajima’s D was consistently negative and statistically insignificant in all datasets, except for Gujarat, where it was positive (0.00096) but statistically insignificant.

#### Positive selection on specific codons of *Pv*puCSP

The Z test of selection dismisses neutrality, favouring purifying selection, but the values lacked statistical significance. The McDonald-Kreitman Test (MKT) was conducted with an out-group, consistently yielding a negative alpha value. In each instance, the neutrality index surpassed one, signifying negative selection, though none of the values achieved statistical significance.

Gene-wide selection using BUSTED shows the evidence of episodic diversifying selection (LRT, p <0.05). It suggests that at least one site on at least one test branch has experienced diversifying selection. A minimal proportion of PvpuCSP sites (1.46%) evolved with a substantial ω value of 536.01.The Evidence-Ratio (constrained and optimized null) value estimated by the program plotted against the sites to check the probable sites that could have evolved under positive selection shows majority of residues lies in the repeat region with high log-likelihood evidence ratio (Fig 4). A minimum of 10 sites within the repeat region exhibited constrained statistics, with an ER value exceeding 10. Notably, one residue at the amino terminal (codon position 33) showed an exceptionally high ER value of 53.47. The comprehensive analysis indicates robust positive selection at specific codon sites within the gene, while the remaining sections of the gene remain highly conserved, characterized by a ω value close to 0

**Figure 4:**
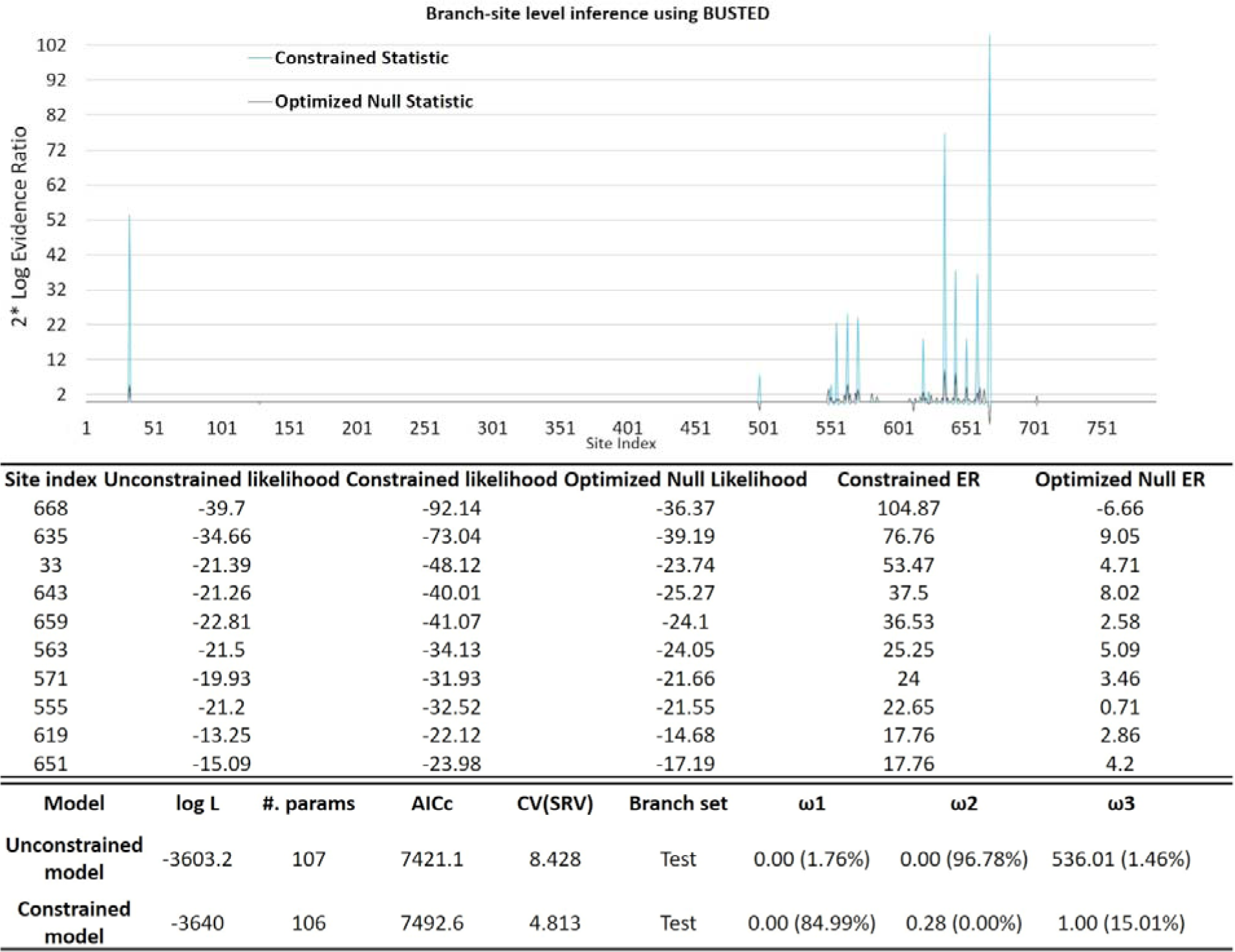
Inference of gene-wide selection using BUSTED. (a)Constrained and optimized null evidence ratio (ER) estimated using BUSTED are plotted across codon sites of *Pv*puCSP. The ER values are scaled to log-likelihood. (b) The codon sites with high LRT (Likelihood Ratio Test) value and evidence ratio, could be the possible sites for selection. (c) The proportion of sites and their corresponding omega rates and the percentage of branches belonging to each category. The constrained model is based on the null hypothesis assuming ω = 1 to obtain background information.

While BUSTED identifies sites under selection, aSBREL provided a combined interpretation at both sites and across genealogy. A total of 83 branches were analysed of which around 75 (90% of branches and 66% of total tree length) branches were evolved with simple model i.e. single ω rate class was allocated to those branches. The other 8 branches (9.6% of branches and 34% of total tree length) were allocated with multiple ω rate classes. aBSREL found evidence of episodic diversifying selection around 2 branches (Node 44 and Node 51) in the phylogeny with statistically significant p-value < 0.05 and likelihood ratio values 27.14 and 18.74, respectively (Fig 5). Node 44 includes the isolates that are exclusively from Delhi population and were collected more recently. The other node which was under selection included two isolates that were from Delhi and Odisha. However, no pattern was observed in the context of population, repeat number and/or arrangement. MEME and FEL identified 12 codon sites (55, 563, 635, 659, 651, 555, 780, 551, 571, 619, 668 and 33) under episodic positive/diversifying selection in *Pv*puCSP with a p-value threshold of 0.1, of which majority of them were present in the repeat region. The 12 codon sites with their corresponding α, β+, LRT score and p-values are mentioned in supplementary table 8. Similarly, FEL which works per site basis and employs a maximum likelihood approach, found evidence of pervasive positive/diversifying selection at 3 sites (634, 643 and 668), with p− value threshold of 0.1.

**Figure 5:**
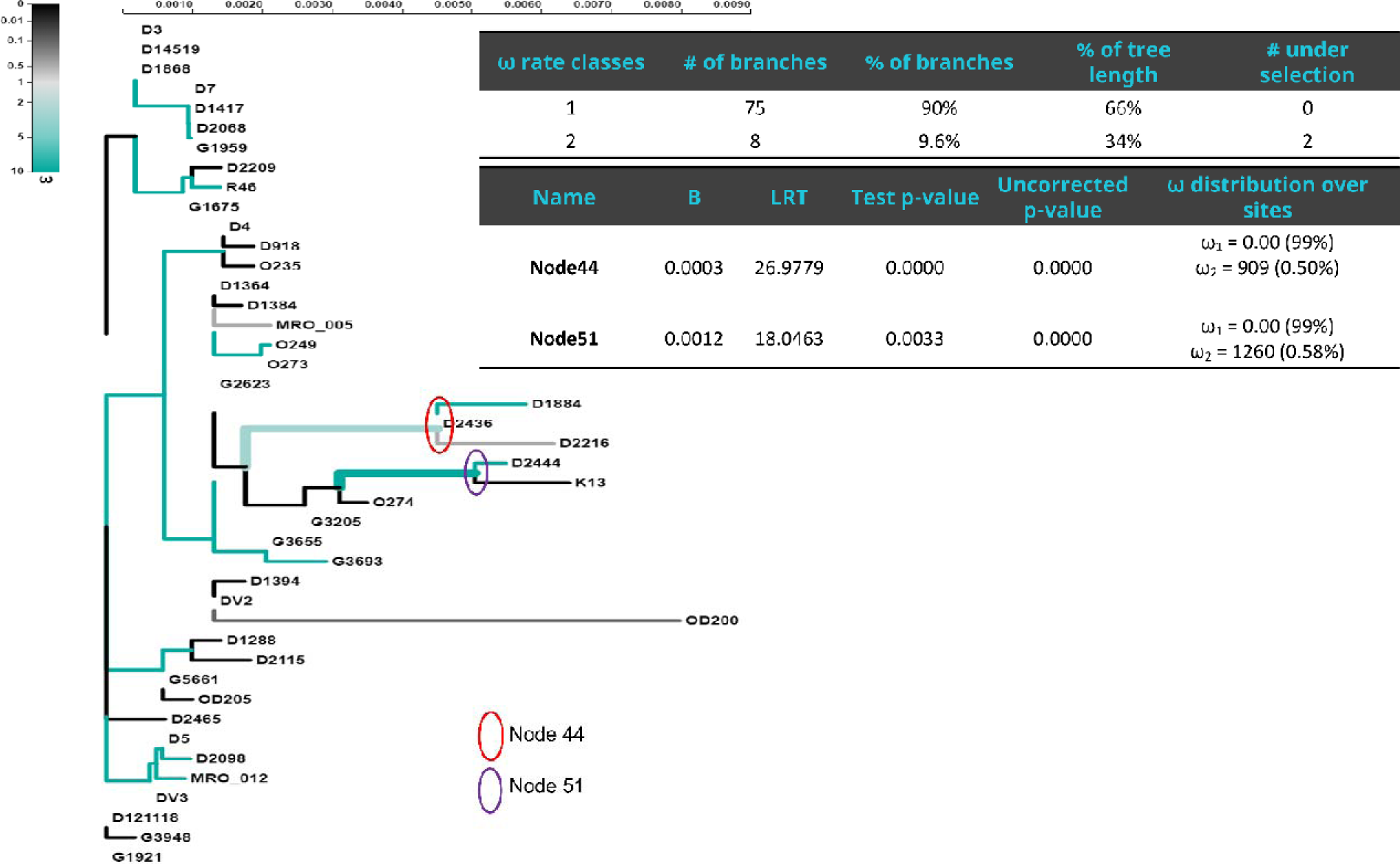
Prediction of branch under selection using the aBSREL method. Phylogenetic tree depicting the fitted aBSREL Adaptive model of *Pv*puCSP. Branches are coloured by their inferred ω distribution. Lineages identified to be under positive selection with statistical significance (p < 0.05) after correction are shown with thick branches.

The strength of selection force also plays a major role and affects the distribution of substitutions. Therefore, the test statistics RELAX was applied to the *Pv*puCSP data to test the strength or intensity of selection along the genealogies, which infers intensity of selection is significant with K= 1.90 (p=0.000, LRT=15.64). Since it was found that strength of selection is intense, it was essential to visualize the degree of linkage disequilibrium and recombination events. Evidence of recombination breakpoint was found at site 1664 with p < 0.05, with left and right-hand sides of the breakpoint had p-a value 0.0002. With one break-point the AICc value of the gene is changed from 7923.53 to 7877.93. Similarly, a single recombination event (Rm) was predicted by DnaSP at the site between 387 and 1643. Linkage disequilibrium with significant Fisher exact test was found at sites between 98 and 2003. Taken together, the recombination breakpoints were predicted to be present at the start and end of the repeat region.

## Discussion

Fixation of a beneficial allele in a population leads to the concurrent fixation of linked non-coding regions (Maynard Smith and Haigh 2008; Schlotterer 2003). Screening such non-functional genomic stretches for decreased diversity and evaluating their linkage with neighbouring functional regions can identify loci that hold both functional and evolutionary significance. For genomes like *P. vivax*, with nearly half of the genes being hypothetical, the mentioned approach helps extract information on significant loci. The study investigates the putative CSP gene (PVX_086150) *in P. vivax*, which lacks functional evidence despite evolutionary significance. Using population genetics, we explore role of selection in shaping the genetic composition of this gene in Indian *P. vivax*.

Tandem repeats in the genome are result from replication errors, unequal crossing over, or duplication events. In *P. falciparum*, indel mutations occur about 10 times more frequently than single nucleotide substitutions, with approximately 9% of genes containing repeat stretches (Hamilton et al. 2017). However, detailed studies on tandem repeats in the *P. vivax* genome are lacking. Mendes et al. found a bias in repeat content, with intracellular parasites having more repeats, and *P. vivax* exhibiting a higher percentage of perfect repeats (Mendes et al. 2013). The evolution of repeats varies based on their location; repeats in noncoding regions may undergo neutral evolution, while those in functional coding regions could be shaped by selection. In human proteins, natural selection influences tandem repeat expansion (Mularoni et al. 2010). Repetitive proteins, like PvpuCSP reported in our previous study, are often intrinsically disordered, showcasing structural plasticity unlike globular proteins (Davies et al. 2017). The variations (Hd: 0.959) in the number of repeats in PvpuCSP in the Indian population suggest repeat accumulations due to trinucleotide slippage and adaptive selection. These changes in repeats likely have functional relevance by providing more surface area for binding proteins targeted for degradation. The absence of repeat units in PvpuCSP orthologs in other Plasmodium species indicates recent and exclusive repeat expansion in *P. vivax*.

PCR amplification and sequencing of PvpuCSP confirmed diverse repeat regions in it. Haplotype analysis, reflecting repeat variation, identified 37 alleles from 71 samples, showcasing the locus’s broad polymorphism. The length differences in *P. vivax* isolates can be easily resolved using gel electrophoresis suggest PvpuCSP as a potential marker for detecting multiplicity of infection (MOI). Differentiating genetically distinct strains is approached at varying levels, employing different markers and techniques (Zhong, Koepfli, et al. 2018). Commonly utilized markers like msp1, msp3alpha, csp, dbp, gam-1, and microsatellites such as Pv3.27, MS16, Pv1.501, and Pv3.502 help estimate MOI in *P. vivax* (Koepfli et al. 2009). PvpuCSP diversity (He: −0.975) is comparable to the highly diverse microsatellite marker MS16 (He: −0.988; Method: Capillary electrophoresis) (Koepfli et al. 2009) and Pvmsp1 (He: −0.969; Method: deep read sequencing) (Zhong, Lo, et al. 2018), currently used for MOI estimation. PCR-RFLP methods face challenges in resolving strains with minimal length variation, particularly in microsatellite markers, risking strain loss. Capillary electrophoresis is crucial for microsatellite-based MOI estimation. PvpuCSP’s 12-nucleotide repeat unit is better resolved in agarose gel electrophoresis, a cost-effective and widely available method. Antigenic markers, commonly used for MOI estimation, may introduce bias due to immune selection pressure, and are not recommended for parasite population structure analysis. PvpuCSP’s non-antigenic nature, single copy, repeat length, and high diversity make it a promising marker for estimating diversity and infection complexity in *P. vivax*. However, further systematic studies comparing PvpuCSP with existing MOI markers are needed to validate its suitability (: Zhong, Lo, et al. 2018).

Investigating the extent of genetic polymorphism in PvpuCSP and exploring the processes that maintain its diversity are crucial for understanding the evolutionary significance of this locus. Despite a common gene structure, with conserved terminals and a variable repeat region, the repeat unit codes for different amino acids (RDNA and RCNA in PvpuCSP, NANP in PfCSP). While PfCSP has a higher repeat unit frequency, PvpuCSP is genetically more diverse in the repeat region due to different allotypes coding for the same motifs. Both CSPs maintain diversity in their repeat regions, with PfCSP exhibiting frequency-dependent immune selection by which both frequent and rare alleles are maintained or favoured by natural selection (Conway 2007). PvpuCSP shows episodic diversifying selection in the repeat region, enhancing amino acid diversity at various phylogenetic levels to improve the adaptation.

Selection pressure can focus on specific sites or a lineage branch, especially in functional loci where most codons face strong structural or functional constraints. Traditional methods, which often rely on average dN/dS, may lack power to detect positive selection in these loci as selection might act on specific sites across different phylogenetic levels rather than the entire locus. Analysing the entire locus may obscure selection signatures. Statistical models capable of detecting selection among lineages or sites are more effective in such scenarios.

BUSTED and MEME, employing a gene-wide test following a branch-site model, detected episodic diversifying selection at specific branches or sites, with statistical significance. Many selected sites, particularly in the repeat region (Figure 5). BUSTED performs a gene-wide test following a branch-site model, revealed episodic diversifying selection at some branches or sites, with statistical significance and many of the sites (codons) are present in the repeat region (Figure 5). aBSREL, a model detecting positive diversifying selection, observed similar episodic diversifying selection at two branches with statistical significance (p<0.05) (Figure 6), aligning with previous analysis.

Recombination breakpoints, purifying selection in the N and C-terminal, and episodic diversifying selection in the repeat region suggest a recombination event, potentially reducing selection strength across the locus. The presence of recombination breakpoints in and around the repeat region, contrasting with high conservation in other gene parts, supports this interpretation. The signal of episodic diversifying selection in the repeat region adds to functional relevance, influenced by recombination dynamics.

Purifying selection dominates the main gene part, affirming its housekeeping nature. The observed variation in the repeat region may enhance protein functionality. Distinguishing between balancing and ongoing positive selection is challenging, as both exhibit similar molecular phenotypes.

Whether the PvpuCSP repeat region variation results from balancing or ongoing positive selection remains uncertain. Given E3 ubiquitin ligases’ role as antimalarial drug targets, understanding PvpuCSP’s genetic diversity and molecular evolution contributes valuable information to potential drug candidates.

## Methods

### Study sites and sample collection

Finger-prick blood samples were collected from individuals across various regions of India with malaria-like symptoms, following passive (patients visiting clinics) and active (direct house survey) approaches (Supplementary Table 1). During active sample collection, bivalent rapid diagnostic test (RDT) kit (FalciVax™, Zephyr Biomedicals, India) was used for prompt identification of *P. falciparum* and *P. vivax*. Positive RDT samples were then used to create dried blood spots (DBS) on 3 mm thick Whatman® filter paper (Whatman, GE Healthcare, UK). The DBS was dried at room temperature, packed in an individual zip-lock pouch with desiccant and transported to ICMR-National Institute of Malaria Research (NIMR) for further processing. The DBS were stored in a −20° C untill further use. The ethics clearance for sample collection was obtained from the Institutional Ethics Committee (IEC) of the ICMR-NIMR, New Delhi, and written informed consents were obtained from patients.

### Sequencing PuCSP gene in mono*-P. vivax* samples

Genomic DNA was isolated from the DBS using QIAmp DNA Minikit (QIAGEN, Germany), following the manufacturer’s instructions. A nested-PCR assay was performed on all the samples to confirm the *Plasmodium* species (Johnston et al. 2006; Siwal et al. 2018), and the samples with mono-infection by *P. vivax* were chosen to amplify and sequence the puCSP gene.

The *Pv*puCSP (*PV*X_086150) gene is a 2.2kb long, single-exon gene, located on the 13^th^ chromosome (chr13:1877708 – 1879927, Sal-I strain) of *P. vivax*. To sequence the whole gene, overlapping primer pairs were designed with first forward and last reverse primers from the flanking region of the gene (Figure 1). Each primer pair was optimized for an optimum annealing temperature (T_a_) and primer concentration (Supplementary Table 2). All the amplification reactions were carried out at a final volume of 25 µl with 10 µM forward and reverse primers, Go Taq® Green Master Mix 2X (Promega Corp., India), genomic DNA template (20 to 30 ng) and nuclease-free water. The amplification reactions were carried out as initial denaturation at 95 ° C for 5 minutes, followed by 35 cycles of (i) denaturation at 95° C for 1 minute, (ii) annealing (60-63° C) for 1 minute, and (iii) extension at 72° C for 1 minute and final elongation at 72° C for 7 minutes.

The amplified products were visualized in 1% agarose gel (Sigma-Aldrich) electrophoresis, with a 1kb DNA marker as reference. *Pv*puCSP’s central repeat region (1636-2000 bp), was amplified with a primer pair outside the repeat region and a sequencing primer within it improved sequencing accuracy (Supplementary Figure 1). The purified PCR products were sequenced with 2x coverage, by capillary electrophoresis in an ABI 3730xl DNA Analyser (Applied Biosystems™), using BigDye™Terminator v3.1 master mix.

Chromatogram data were manually reviewed and analysed using DNAStar Lasergene’s SeqMan module (Madison, WI, USA) to eliminate base calling errors (Q score <20) and discard sub-threshold stretches. The fragments were assembled with “match size: 12 at a “minimum match percentage of 80” and “minimum sequence length of 100” to generate the complete gene. Unassembled fragments were rejected and re-sequenced. Chromatograms were scrutinized for single peaks at each nucleotide position, indicating a singular infection. Multiple Alignment Fast Fourier Transform (MAFFT) with *L-ins-i* parameter was used for multiple sequence alignment (Katoh et al. 2005). The gaps created during alignment were considered as InDel during polymorphism.

### Analysis of *Pv*puCSP central repeat region

Our prior research showed the presence of a variable tetra-peptide repeat unit (R-D-N-A) at the central region of the locus, among isolates (Dash et al. 2019). Therefore, sequenced loci were screened for repeats using Tandem Repeats Finder (TRF) (Benson 1999). The repeat units are grouped into repeat allotypes (RATs) and protein repeat motifs (PRM).

Alleles were defined based on RAT arrangement in the repeat region. Allelic richness was estimated using Allelic Diversity analyZEr (ADZE), which uses a rarefaction approach to correct for biasedness from uneven sample sizes among populations (Szpiech, Jakobsson, and Rosenberg 2008). Summary statistics (haplotype diversity, expected heterozygosity) and allele distribution (unique and shared alleles) were determined for each population using a “haploid” model in the GenAlex program (Peakall and Smouse 2012).

### Analysis of the *Pv*puCSP based on SNP distribution

The repeat regions were removed before estimating genetic diversity and summary statistics based on SNP distribution, using DnaSP v6. The locus was analysed as a whole and by grouping them into N-terminal (Alignment position:1-1635 bp), central repeat region (1636-1995 bp), and C-terminal (1996-2376 bp). A similar approach was followed to estimate the genetic diversity of the complete *Pv*puCSP using GenAlex and allelic richness approach.

### Tests of neutrality and recombination

The neutral theory of molecular evolution was evaluated using various frequency-based models, using DnaSP (Rozas et al. 2017). The codon-based Z-test, with Jukes-Cantor correction and McDonald-Krietman (MK) test (McDonald and Kreitman 1991) was implemented using MEGA X (Kumar et al. 2018). *P. cynomolgi* was used as an outgroup to estimate divergence. The mutation-based tests average omega (dN/dS) across branches and codon sites, reducing their ability to detect localised or transient selection (Murrell et al. 2015). Branch-site model tests (e.g., aBSREL and BUSTED) on *Pv*puCSP assessed stochastic selection impact across codon-sites and branches using Data Monkey server (Murrell et al. 2015; Weaver et al. 2018). Degree of linkage disequilibrium (LD), recombination events (Rm), and recombination sites were assessed with DnaSP V6. Expected LD was determined using the correlation coefficient with Bonferroni correction, and significance was tested with Fisher’s exact test (Hill & Robertson 1968). Recombination breakpoints were identified using GARD and SBP programs on the Data Monkey server (Pond et al. 2006).

## Acknowledgment

We thank all the donors who have given their blood samples to carry out the study. We also thank director NIMR for providing necessary research facilities and moral support during the study.

Thanks are also owing to all the lab members for their helpful comments and suggestions while preparing the manuscript.

## Author contribution

MD collected the samples, performed the experiments, generated and analysed the data, wrote the manuscript, and produced the figures and tables. VP designed the study, critically reviewed the manuscript. AD conceptualized the study and critically reviewed the manuscript. AS conceptualized and designed the study, planned the experiment and analysis, critically reviewed and edited the manuscript. All authors agreed to the final version of the manuscript for publication.

## References

Barton, N. H. 2000. “Genetic Hitchhiking.” Philosophical Transactions of the Royal Society of London. Series B, Biological Sciences 355(1403):1553–62.

Benson, Gary. 1999. “Tandem Repeats Finder:A Program to Analyze DNA Sequences.” Nucleic Acids Res 27(2):573–80.

Carlton, Jane M., John H. Adams, Joana C. Silva, Shelby L. Bidwell, Hernan Lorenzi, Elisabet Caler, Jonathan Crabtree, Samuel V Angiuoli, Emilio F. Merino, Paolo Amedeo, Qin Cheng, Richard M. R. Coulson, Brendan S. Crabb, Hernando A. Del Portillo, Kobby Essien, Tamara V Feldblyum, Carmen Fernandez-Becerra, Paul R. Gilson, Amy H. Gueye, Xiang Guo, Simon Kang’a, Taco W. A. Kooij, Michael Korsinczky, Esmeralda V. S. Meyer, Vish Nene, Ian Paulsen, Owen White, Stuart A. Ralph, Qinghu Ren, Tobias J. Sargeant, Steven L. Salzberg, Christian J. Stoeckert, Steven A. Sullivan, Marcio M. Yamamoto, Stephen L. Hoffman, Jennifer R. Wortman, Malcolm J. Gardner, Mary R. Galinski, John W. Barnwell, and Claire M. Fraser-Liggett. 2008. “Comparative Genomics of the Neglected Human Malaria Parasite Plasmodium Vivax.” Nature 455(7214):757–63.

Carlton, Jane M., Ananias A. Escalante, Daniel Neafsey, and Sarah K. Volkman. 2008. “Comparative Evolutionary Genomics of Human Malaria Parasites.” Trends in Parasitology 24(12):545–50.

Conway, David J. 2007. “Molecular Epidemiology of Malaria.” Clinical Microbiology Reviews 20(1):188–204.

Das, Aparup. 2015. “The Distinctive Features of Indian Malaria Parasites.” Trends in Parasitology 31(3):83–86.

Dash, M., V. Pande, and A. Sinha. 2019. “Putative Circumsporozoite Protein (CSP) of Plasmodium Vivax Is Considerably Distinct from the Well-Known CSP and Plays a Role in the Protein Ubiquitination Pathway.” Gene 721S(100024).

Davies, Heledd M., Stephanie D. Nofal, Emilia J. McLaughlin, and Andrew R. Osborne. 2017. “Repetitive Sequences in Malaria Parasite Proteins.” FEMS Microbiology Reviews 41(6):923–40.

Day, K. P. and K. Marsh. 1991. “Naturally Acquired Immunity to Plasmodium Falciparum.” Immunology Today 12(3):A68–71.

Fay, Justin C. 2011. “Weighing the Evidence for Adaptation at the Molecular Level.” Trends in Genetics 27(9):343–49.

Fay, Justin C. and Chung I. Wu. 2000. “Hitchhiking under Positive Darwinian Selection.” Genetics 155(3):1405–13.

Ferreira, Marcelo U., Mônica da Silva Nunes, and Gerhard Wunderlich. 2004. “Antigenic Diversity and Immune Evasion by Malaria Parasites.” Clinical and Diagnostic Laboratory Immunology 11(6):987–95.

Fu, Y. X. and W. H. Li. 1993. “Statistical Tests of Neutrality of Mutations.” Genetics 133(3):693–709.

Fu, Yun Xin. 1997. “Statistical Tests of Neutrality of Mutations against Population Growth, Hitchhiking and Background Selection.” Genetics 147(2):915–25.

Gupta, Bhavna, Aditya P. Dash, Nalini Shrivastava, and Aparup Das. 2010a. “Single Nucleotide Polymorphisms, Putatively Neutral DNA Markers and Population Genetic Parameters in Indian Plasmodium Vivax Isolates.” Parasitology 137:1721–30.

Gupta, Bhavna, Aditya P. Dash, Nalini Shrivastava, and Aparup Das. 2010b. “Single Nucleotide Polymorphisms, Putatively Neutral DNA Markers and Population Genetic Parameters in Indian Plasmodium Vivax Isolates.” Parasitology 137(12):1721–30.

Gupta, Bhavna, Nalini Srivastava, and Aparup Das. 2012. “Inferring the Evolutionary History of Indian Plasmodium Vivax from Population Genetic Analyses of Multilocus Nuclear DNA Fragments.” Molecular Ecology 21(7):1597–1616.

Habtamu, Kassahun, Beyene Petros, and Guiyun Yan. 2022. “Plasmodium Vivax: The Potential Obstacles It Presents to Malaria Elimination and Eradication.” Tropical Diseases, Travel Medicine and Vaccines 8(1):27.

Hamilton, William L., Antoine Claessens, Thomas D. Otto, Mihir Kekre, Rick M. Fairhurst, Julian C. Rayner, and Dominic Kwiatkowski. 2017. “Extreme Mutation Bias and High AT Content in Plasmodium Falciparum.” Nucleic Acids Research 45(4):1889–1901.

Johnston, Stephanie P., Norman J. Pieniazek, Maniphet V. Xayavong, Susan B. Slemenda, Patricia P. Wilkins, and Alexandre J. Da Silva. 2006. “PCR as a Confirmatory Technique for Laboratory Diagnosis of Malaria.” Journal of Clinical Microbiology 44(3):1087–89.

Katoh, Kazutaka, Kei Ichi Kuma, Hiroyuki Toh, and Takashi Miyata. 2005. “MAFFT Version 5: Improvement in Accuracy of Multiple Sequence Alignment.” Nucleic Acids Research 33(2):511–18.

Koepfli, Cristian, Ivo Mueller, Jutta Marfurt, Mary Goroti, Albert Sie, Olive Oa, Blaise Genton, Hans Peter Beck, and Ingrid Felger. 2009. “Evaluation of Plasmodium Vivax Genotyping Markers for Molecular Monitoring in Clinical Trials.” Journal of Infectious Diseases 199(7):1074–80.

Kumar, Sudhir, Glen Stecher, Michael Li, Christina Knyaz, and Koichiro Tamura. 2018. “MEGA X: Molecular Evolutionary Genetics Analysis across Computing Platforms.” Molecular Biology and Evolution 35(6):1547–49.

Loy, E. Dorothy, Weimin Liu, Yingying Li, H. Gerald LEarn, J. Lindsey Plenderleith, Sesh a Sundararaman, PAul M. Sharp, and Beatrice H. Hahn. 2017. “Out of Africa: Origins and Evolution of the Human Malaria Parasites Plasmodium Falciparum and Plasmodium Vivax Dorothy.” International Journal for Parasitology 47:87–97.

Maynard Smith, John and John Haigh. 2008. “The Hitch-Hiking Effect of a Favourable Gene.” Genetics Research 89(5–6):391–403.

McDonald, J. H. and M. Kreitman. 1991. “Adaptive Protein Evolution at the Adh Locus in Drosophila.” Nature 351(6328):652–54.

Mendes, T. A. O., F. P. Lobo, T. S. Rodrigues, G. F. Rodrigues-Luiz, W. D. daRocha, R. T. Fujiwara, S. M. R. Teixeira, and D. C. Bartholomeu. 2013. “Repeat-Enriched Proteins Are Related to Host Cell Invasion and Immune Evasion in Parasitic Protozoa.” Molecular Biology and Evolution 30(4):951–63.

Messer, Philipp W. and Dmitri A. Petrov. 2013. “Frequent Adaptation and the McDonald – Kreitman Test.” 2013.

Mularoni, Loris, Alice Ledda, Macarena Toll-Riera, and M. Mar Albà. 2010. “Natural Selection Drives the Accumulation of Amino Acid Tandem Repeats in Human Proteins.” Genome Research 20(6):745–54.

Murrell, Ben, Steven Weaver, Martin D. Smith, Joel O. Wertheim, Sasha Murrell, Anthony Aylward, Kemal Eren, Tristan Pollner, Darren P. Martin, Davey M. Smith, Konrad Scheffler, and Sergei L. Kosakovsky Pond. 2015. “Gene-Wide Identification of Episodic Selection.” Molecular Biology and Evolution 32(5):1365–71.

Peakall, Rod and Peter E. Smouse. 2012. “GenAlEx 6.5: Genetic Analysis in Excel. Population Genetic Software for Teaching and Research--an Update.” Bioinformatics (Oxford, England) 28(19):2537–39.

Pond, Sergei L. Kosakovsk., David Posada, Michael B. Gravenor, Christopher H. Woelk, and Simon D. W. Frost. 2006. “Automated Phylogenetic Detection of Recombination Using a Genetic Algorithm.” Molecular Biology and Evolution 23(10):1891–1901.

Rozas, Julio, Albert Ferrer-Mata, Juan Carlos Sánchez-DelBarrio, Sara Guirao-Rico, Pablo Librado, Sebastián E. Ramos-Onsins, and Alejandro Sánchez-Gracia. 2017. “DnaSP 6: DNA Sequence Polymorphism Analysis of Large Data Sets.” Molecular Biology and Evolution 34(12):3299–3302.

Schlotterer, Christian. 2003. “Hitchhiking Mapping – Functional Genomics from the Population Genetics Perspective.” Trends in Genetics 19(1):32–38.

Siwal, Nisha, Upasana Shyamsunder Singh, Manoswini Dash, Sonalika Kar, Swati Rani, Charu Rawal, Rajkumar Singh, Anupkumar R. Anvikar, Veena Pande, and Aparup Das. 2018. “Malaria Diagnosis by PCR Revealed Differential Distribution of Mono and Mixed Species Infections by Plasmodium Falciparum and *P. vivax* in India.” PloS One 13(3):e0193046.

Szpiech, Zachary A., Mattias Jakobsson, and Noah A. Rosenberg. 2008. “ADZE: A Rarefaction Approach for Counting Alleles Private to Combinations of Populations.” Bioinformatics (Oxford, England) 24(21):2498–2504.

Tang, Haibao, John E. Bowers, Xiyin Wang, Ray Ming, Maqsudul Alam, and Andrew H. Paterson. 2008. “Synteny and Collinearity in Plant Genomes.” Science (New York, N.Y.) 320(5875):486–88.

Weaver, Steven, Stephen D. Shank, Stephanie J. Spielman, Michael Li, Spencer V Muse, and Sergei L. Kosakovsky Pond. 2018. “Datamonkey 2.0: A Modern Web Application for Characterizing Selective and Other Evolutionary Processes.” Molecular Biology and Evolution 35(3):773–77.

Zhong, Daibin, Cristian Koepfli, Liwang Cui, and Guiyun Yan. 2018. “Molecular Approaches to Determine the Multiplicity of Plasmodium Infections.” Malaria Journal 17(1):1–9.

Zhong, Daibin, Eugenia Lo, Xiaoming Wang, Delenasaw Yewhalaw, Guofa Zhou, Harrysone E. Atieli, Andrew Githeko, Elizabeth Hemming-Schroeder, Ming Chieh Lee, Yaw Afrane, and Guiyun Yan. 2018. “Multiplicity and Molecular Epidemiology of Plasmodium Vivax and Plasmodium Falciparum Infections in East Africa.” Malaria Journal 17(1):1–14.

